# The proteomic toolbox for identification, quantification, and characterization of polyclonal antibodies

**DOI:** 10.1101/2023.10.27.564451

**Authors:** Weize Tang, Andrei P. Drabovich

## Abstract

Recent advances in proteomics and mass spectrometry facilitated the in-depth characterization of monoclonal antibodies and enabled innovative approaches for the quantification of polyclonal antibodies generated against numerous antigens. Human respiratory syncytial virus (RSV) is a contagious respiratory pathogen often manifested as a common cold infection in adults and more serious symptoms in infants and the elderly population. Here, we used a reference IgG1κ monoclonal antibody NISTmAb 8671 and its affinity interaction with an RSV fusion glycoprotein F as a model to develop the proteomic toolbox for identification, quantification, and characterization of polyclonal antibodies. Our toolbox integrated a variety of proteomic and mass spectrometry approaches for accurate mass measurements of antibody fragments, antibody digestion with the complimentary proteases (trypsin, asparaginase, and proalanase), immunoaffinity enrichments, and bottom-up or middle-down proteomics. We measured absolute concentrations of anti-RSV antibody isotypes and subclasses in 69 serum samples of healthy individuals and revealed IgG1 (2,580 ng/mL), IgA1 (280 ng/mL), and IgM (180 ng/mL) as the most abundant isotypes. Interestingly, we also identified the presence of IgG2 (74 ng/ml), IgG4 (4.9 ng/mL) and IgA2 (5.5 ng/mL) antibodies. Interactome measurements detected the consistent co-precipitation of C1q complement complexes. Repertoire profiling of the variable regions of polyclonal antibodies revealed the frequent use of IGHV3 subgroup genes, while IGHV5-51 was the most abundant single gene of the anti-RSV polyclonal antibody response. The presented toolbox will facilitate the in-depth characterization of polyclonal antibodies and pave the way to quantitative approaches in serological studies and precision immunology.

## INTRODUCTION

Evaluation of individual and population immunity often relies on the detection of antigen- or pathogen-specific antibodies in biological fluids, such as blood serum. Enzyme-linked immunosorbent assay (ELISA) is a conventional tool for serological testing and is used clinically to provide diagnosis of numerous diseases. Indirect ELISA, however, has limitations such as semi-quantitative measurements, lack of standardization, cross-reactivity, and non-specific binding. Innovative analytical techniques such as proteomics and mass spectrometry (MS) promise to resolve some of those limitations. Here, we present a proteomic toolbox for the identification, quantification, and characterization of polyclonal antibodies. The proposed ImmunoAffinity-Mass Spectrometry (IA-MS) approaches could overcome the challenges of indirect ELISA and provide additional knowledge on antibody diversity, abundance, and cooperation in immune response.

To demonstrate the advantages of proteomics and IA-MS for comprehensive serological studies, we selected as a model the polyclonal antibodies generated against the respiratory syncytial virus (RSV). Human RSV is a common virus that causes infections of the respiratory tract in infants and numerous reinfections in later life, with its polyclonal antibodies detected in the blood serum of nearly all adults^1^. RSV results in over 3 million hospitalizations and up to 199,000 childhood deaths worldwide^2^. Infants typically obtain anti-RSV antibodies via placental transfer from mothers and become protected for the first few months of their life. While nearly all adults have circulating serum anti-RSV antibodies due to previous reinfections, antibody levels decrease in the elderly population and immunocompromised patients. Measurement of the circulating levels of anti-RSV antibodies may provide prognostic information for the severity of RSV infections in infants, the elderly, and immunocompromised patients.

Previous studies identified pre-fusion glycoprotein F (preF) as the most immunogenic RSV protein and revealed the increase of the titer of anti-preF IgG1, IgG2 and IgG3 antibody subclasses after RSV infections^3^. Indirect ELISA studies with a variety of calibrators (palivizumab, NISTmAb, human reference serum, etc.) revealed vastly inconsistent concentrations of anti-preF IgG antibodies in convalescent adult sera ranging from 0.3 µg/ml^1^ to 2-4 µg/ml^4,5^ to 30 µg/mL^6^. Two order magnitude differences of the reported average concentrations revealed the challenges with assay standardization and would justify the need for independent gold standard assays.

The diversity of polyclonal antibodies generated against specific antigens could now be approximated with the diversity of the B cell receptor (BCR) repertoires measured by single-cell RNA sequencing^7^. Sequencing of memory B cells or plasmablasts isolated from the human peripheral blood mononuclear cells (PBMCs) by the flow sorting techniques is a common approach to identify BCR repertoires, complementarity-determining region 3 (CDR3) hypervariable sequences, and the frequency of usage of IGHV and IGHJ germline genes in the BCR repertoires^4^. Single-cell BRC sequencing is a powerful approach but has certain limitations: (i) memory B cells and plasmablasts isolated from PBMCs are physically and functionally separated from the antibody-secreting long-lived plasma B cells^8,9^ which reside in bone marrow and can hardly be obtained from healthy donors; (ii) long-lived plasma B cells may have heavily mutated BCR repertoires and higher affinities of antibodies as compared to memory B cells^10,11^; (iii) BRC sequencing cannot predict the final concentration, affinity, and diversity of antibody clones present in circulation; (iv) BCR sequencing relies on RT-PCR amplification and is prone to additional mutations and errors which can hardly be ruled out in CDR regions due to the lack of templates (the challenges of *de novo* versus template-based assembly); (v) BCR sequencing studies often report incomplete or truncated sequences missing information for ∼15 amino acids at the N-terminus. Novel approaches for the direct evaluation of the abundance, affinity, and repertoire diversity of polyclonal antibodies could complement and advance the BCR RNAseq studies.

We believe that the proposed proteomic and ImmunoAffinity-Mass Spectrometry (IA-MS) toolbox will facilitate: (i) direct measurements of absolute concentrations of antigen-specific polyclonal antibodies; (ii) differential quantification of immunoglobulin isotypes and subclasses; (iii) increased VH sequence coverage using the alternative proteases; (iv) matching of V and L chains of antibody clones by accurate measurements of Fab, Fc and L chain masses; (v) measurement of the frequency and relative abundance of IGHV usage; and (vi) identification of Fc interactome. The presented toolbox will facilitate quantitative approaches in serological diagnostics and precision immunology, and enable in-depth characterization of polyclonal antibodies.

## EXPERIMENTAL METHODS

### Reagents and samples

Recombinant RSV fusion glycoprotein F (preF) extracellular domain (529 aa; #11049-V08B) was obtained from Sino Biological. Synthetic stable isotope-labeled peptide standards were provided by JPT Peptide Technologies. Dithiothreitol and iodoacetamide were obtained from Sigma. Tris(2-carboxyethyl)phosphine hydrochloride (TCEP; #20490) was obtained from ThermoFisher Scientific. NIST monoclonal antibody reference material (NISTmAb 8671) was obtained from the National Institute of Standards and Technology^12^. MS-grade ProAlanase (#VA2171), asparaginase (#VVA1160), and Trypsin (#V5111) were obtained from Promega. Cysteine protease IgdE (FabALACTICA; #A0-AG1-020) was obtained from Genovis. The use of archived deidentified serum samples previously collected from healthy individuals was approved by the University of Alberta (#Pro00104098) and the Health Research Ethics Board of Alberta (#HREBA.CC-22-0056).

### Generation of Fab fragments

NISTmAb monoclonal antibody (300 µg) was digested with 30 units of IgdE to generate Fab and Fc fragments. Fc/2, Fd, and L chain fragments were generated by reduction of disulfide bonds with TCEP.

### Native protein LC-MS

NISTmAB fragments (40 ng) were separated using EASY-nLC II nanoLC (Thermo Scientific) with a Polystyrene Divinylbenzene EASY-Spray Capillary HPLC Column (150 μm x 150 mm, 4 μm particles; ES907) over 20-40% acetonitrile gradient at 1 μl/min flow rate. NISTmAB fragments were ionized with an EASY-Spray source, and analyzed with Orbitrap Elite mass spectrometer (Thermo Scientific). Profile MS1 scans (1000-2500 m/z; 15K resolution) included 10 microscans, 2.4 kV source voltage, capillary temperature 290°C, and 1e6 FTMS full AGC target. Raw MS files were loaded to UniDec GIU (v6.0.3) software and UniChrom module to calculate the intact masses of Fab, Fc/2, Fd, and L chain fragments.

### Immunoprecipitation

High-binding microplates were coated overnight with recombinant RSV preF protein (0.5 µg/well), blocked with 2% bovine serum albumin (BSA), and washed. Serum (20 μL) was diluted with 0.1% BSA, incubated for 2 hours, and washed with 50 mM ammonium bicarbonate.

### Proteomic sample preparation

Following 2 h on-plate incubation and washing, enriched antibodies were reduced with dithiothreitol, alkylated with iodoacetamide, and digested on the same plate with trypsin (0.25 µg/well), or ProAlanase (0.25 µg/well), or asparaginase (0.25 µg/well). SpikeTides_TQL peptide internal standards (300 fmol/well) were spiked before digestion. Tryptic peptides were concentrated with C18 microextraction.

### Shotgun LC-MS/MS

Peptides were separated using EASY-nLC II nanoLC (Thermo Scientific) with an EASY-Spray Capillary HPLC Column (75 μm x 150 mm, 3 μm C18 particles; ES900) or an µPAC HPLC Column (200 cm, Thermo Scientific, #COLNANO200G1B) over 5-40% acetonitrile gradient at 300 nl/min flow rate. Peptides were ionized with an EASY-Spray source and analyzed with Orbitrap Elite mass spectrometer (Thermo Scientific). Profile MS1 scans (303-1450 m/z; 60K resolution; lock mass 445.120025 m/z) were followed by top 15 Orbitrap MS/MS scans (200-1600 m/z) after HCD fragmentation at normalized CE of 30. Dynamic exclusion included 35 s and repeat count 1. Tuning included 2.4 kV source voltage, capillary temperature 290°C, and 1e6 FTMS full AGC target, 100 ms Full and MSn ion time. Raw files were searched with MaxQuant v2.4.0 and UniProt human canonical proteome database (20386 entries, July 7, 2022) with the additional sequences of the 44 most frequent human germline IGHV genes and alleles (>0.5% frequency in the population), 7,272 IGHV sequences of the human antibodies derived by single-cell RNA sequencing of human B cells, and sequences of NISTmAb, RSV preF protein, BSA, trypsin, proalanase and asparaginase. Modifications included constant cysteine carbamidomethylation and variable methionine oxidation, glutamine and asparagine deamidation, and peptide N-term pyroglutamic acid formation for glutamine and glutamic acid. Up to four missed cleavages were searched for ProAlanase and asparaginase, and three missed cleavages for trypsin. Volcano plots were prepared with Perseus software (v2.0.11). Draw Map web tool was used for sequence coverage visualization^13^.

### Differential quantification of anti-RSV immunoglobulin isotypes and subclasses by LC-MS/MS

Skyline software (v 23.0.9) was used to extract MS1 peaks (MS1 filtering at 60K resolving power at 400m/z) for the light indigenous and heavy isotope-labeled peptides which represented individual isotypes and subclasses, as previously reported^14^. The combined area of three precursor states (M, M+1, M+2) was used to calculate the light-to-heavy (L/H) ratio. Serum volume (20 ul), amount of heavy peptide internal standards (300 fmol/well), and L/H ratios were used to calculate the absolute concentration of each isotype and subclass.

## RESULTS AND DISCUSSION

### Characterization of NISTmAb 8671 and its fragments by native MS

The NIST monoclonal antibody reference material 8671 (NISTmAb 8671)^12^ has been developed and offered as a certified reagent to evaluate and standardize analytical methods to measure monoclonal antibodies^15^. NISTmAb is a recombinant humanized IgG1 kappa antibody (∼150 kDa homodimer of two identical light and heavy chains). The heavy chain has several post-translational modifications including N-terminal pyroglutamination, C-terminal lysine clipping, and the variable glycosylation pattern^15^. Large molecular weight and variable post-translational modifications, such as glycosylation, impede the accurate mass measurements of intact NISTmAb 8671 using native MS. In this work, we demonstrated that NISTmAb cleavage by IgdE protease will generate Fab fragments devoid of variable glycosylation and C-terminal lysine clipping and will facilitate straightforward measurements of accurate masses of Fab, Fd, and L chain fragments (**Fig. 1**). Determination of the accurate masses may enable straightforward VH-VL chain matching for the mixtures of monoclonal antibodies, and potentially simple mixtures of polyclonal antibodies.

**Fig. 1.**
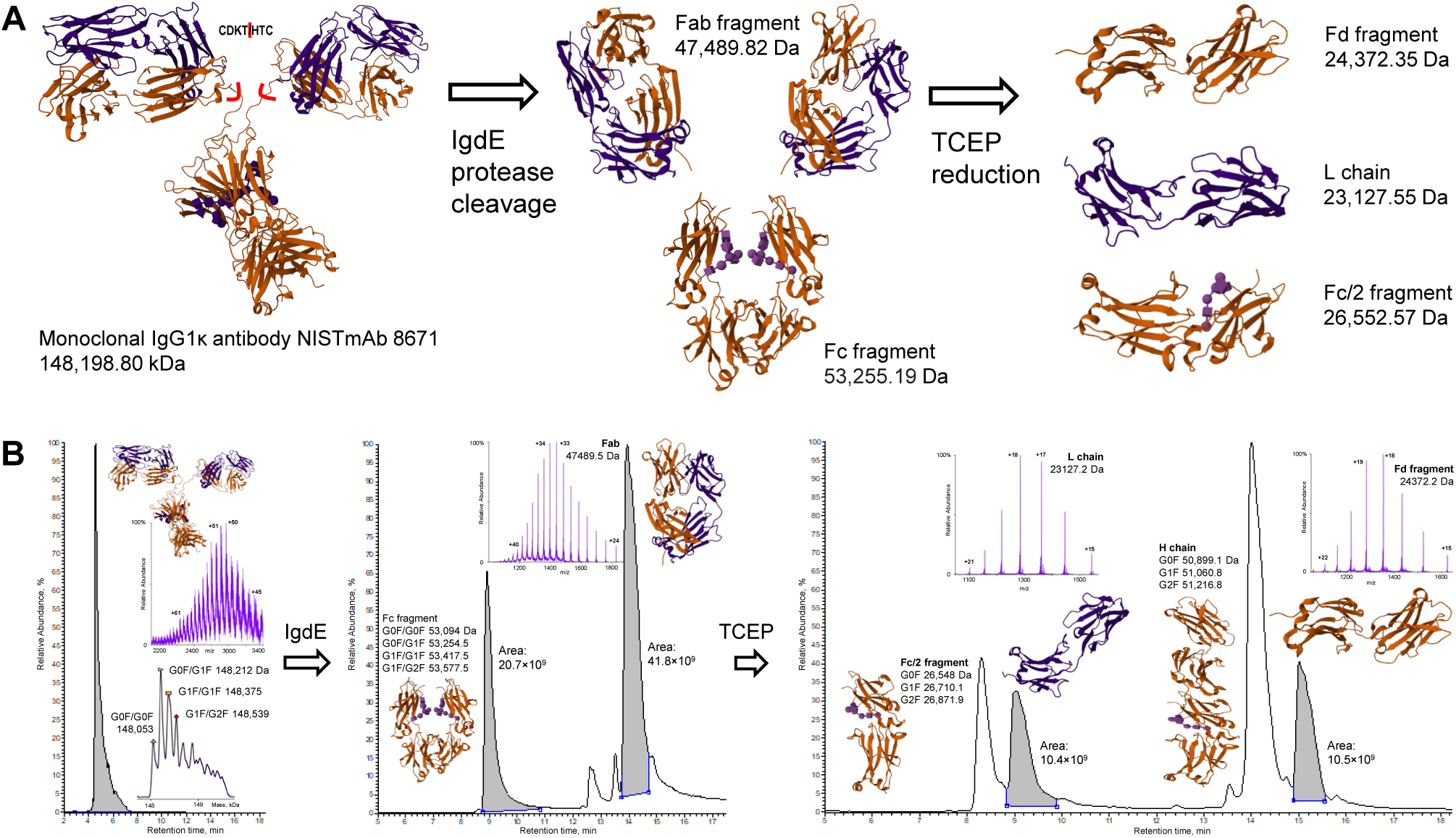
Characterisation of NISTmAb. **(A)** An experimental approach to generate Fab, Fc, Fd, Fc/2, and L chain fragments. **(B)** Native MS to measure accurate masses of Fab, Fd, and L fragments. The schematic structure of a monoclonal antibody was generated with a PDB 1IGT structure.

### Sequence coverage of NISTmAb following digestion with trypsin, asparaginase, and proalanase

We previously demonstrated the numerous applications of shotgun and targeted mass spectrometry to identify and quantify human proteins^16–30^, discover protein biomarkers^31–44^, and investigate the functional roles of human proteins^45–54^. While our previous studies nearly exclusively utilized trypsin-based proteomic approaches, the availability of novel high-quality proteases extended the variety of experimental tools available for proteomic studies. The use of multiple and complementary proteases to characterize and sequence monoclonal antibodies is now an established approach^55–57^. Proalanase is a novel and promising protease that cleaves peptide sequences after proline and alanine^58^. Here, we evaluated proalanase as a tool for NISTmAb sequencing by nanoflowLC-MS/MS with higher-energy collisional dissociation (HCD) fragmentation (**Fig. 2**). The sequence coverage of trypsin-generated peptides was superior (>80%), while asparaginase and proalanase revealed ∼60% sequence coverage. Interestingly, proalanase provided sequencing of some regions not covered by trypsin or asparaginase. The addition of proalanase increased trypsin sequence coverage by 12% (83 to 95%), while the addition of asparaginase provided only a 4% increase. Even though three proteases provided a combined coverage of ∼97% (**Fig. 2**), the contribution of asparaginase was less substantial. While our results need to be confirmed with the additional monoclonal antibodies, we believe that proalanase could emerge as the protease of choice to complement trypsin for the in-depth sequencing of antibodies.

**Fig. 2.**
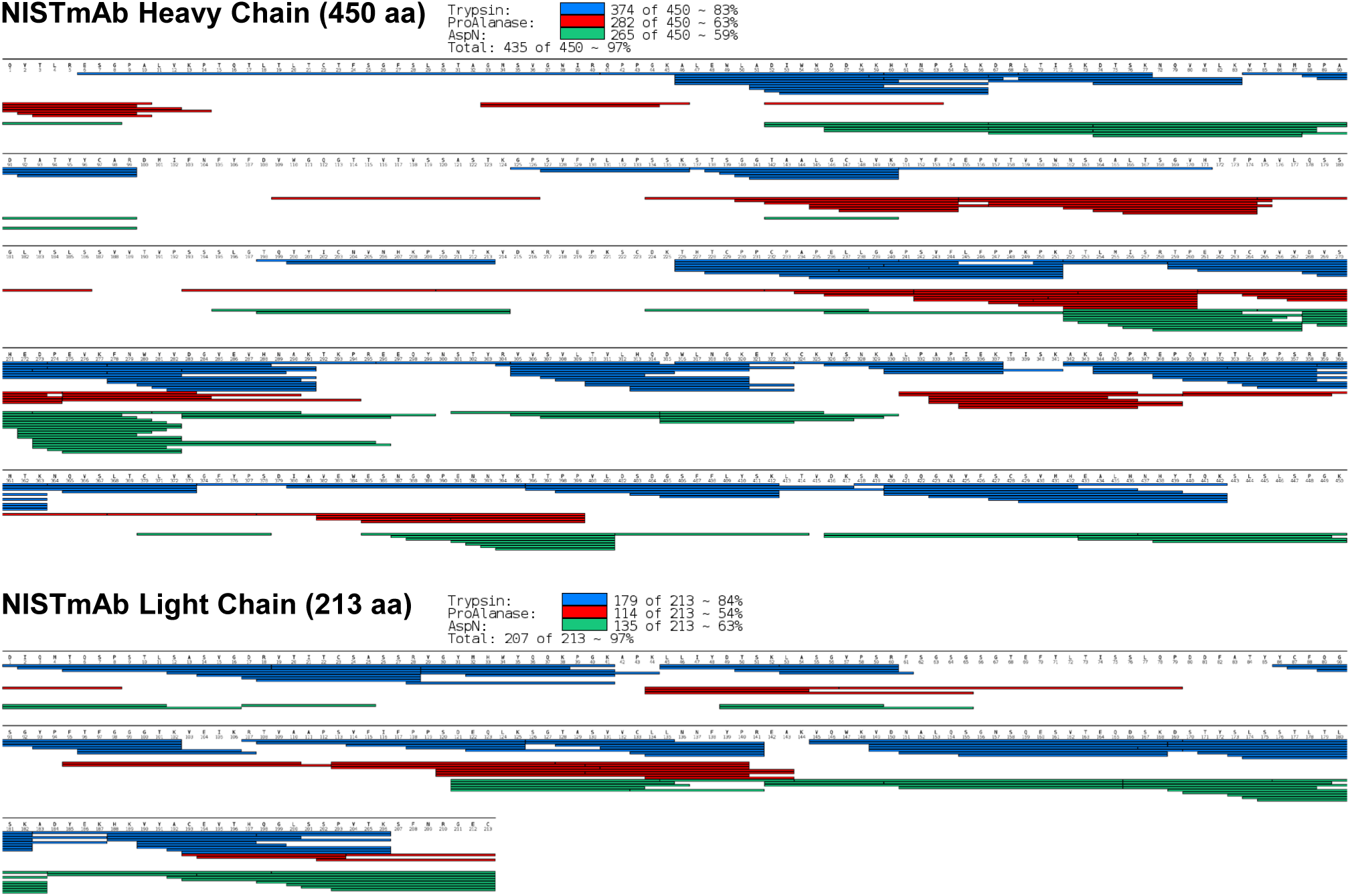
Sequence coverage of NISTmAb 8671 heavy and light chains following digestion with trypsin, asparaginase, and proalanase.

### Development of IA-MS assay to measure the interaction of NISTmAb with RSV preF protein

Our motivation to develop a model IA-MS assay was justified by its future use to map epitopes of monoclonal antibodies and characterize the endogenous polyclonal antibodies. Interaction of NISTmAb with preF protein was chosen as an excellent model due to: (i) high affinity of binding (K_d_ = 4 nM); (ii) a known epitope NSELLSLINDMPITNDQKKLMSNN^59^; (iii) high affinity of binding to the linear peptide epitope (66 nM)^60^; (iv) availability and affordability of a high-quality purified NISTmAb standard; and (v) availability of several monoclonal antibodies with different affinities (palivizumab, motavizumab, and nirsevimab^61^) which could be used as a model to separate antibody mixtures by their affinities, as we previously demonstrated for combinatorial DNA libraries^62–71^. We demonstrated that NISTmAb 8671 was efficiently enriched with preF coated onto microplates (**Fig. 3**). NanoLC-MS/MS with label-free MS1 quantification demonstrated 640-fold enrichment of NISTmAb 8671 with preF, as compared to PBS controls (**Fig. 3**).

**Fig. 3.**
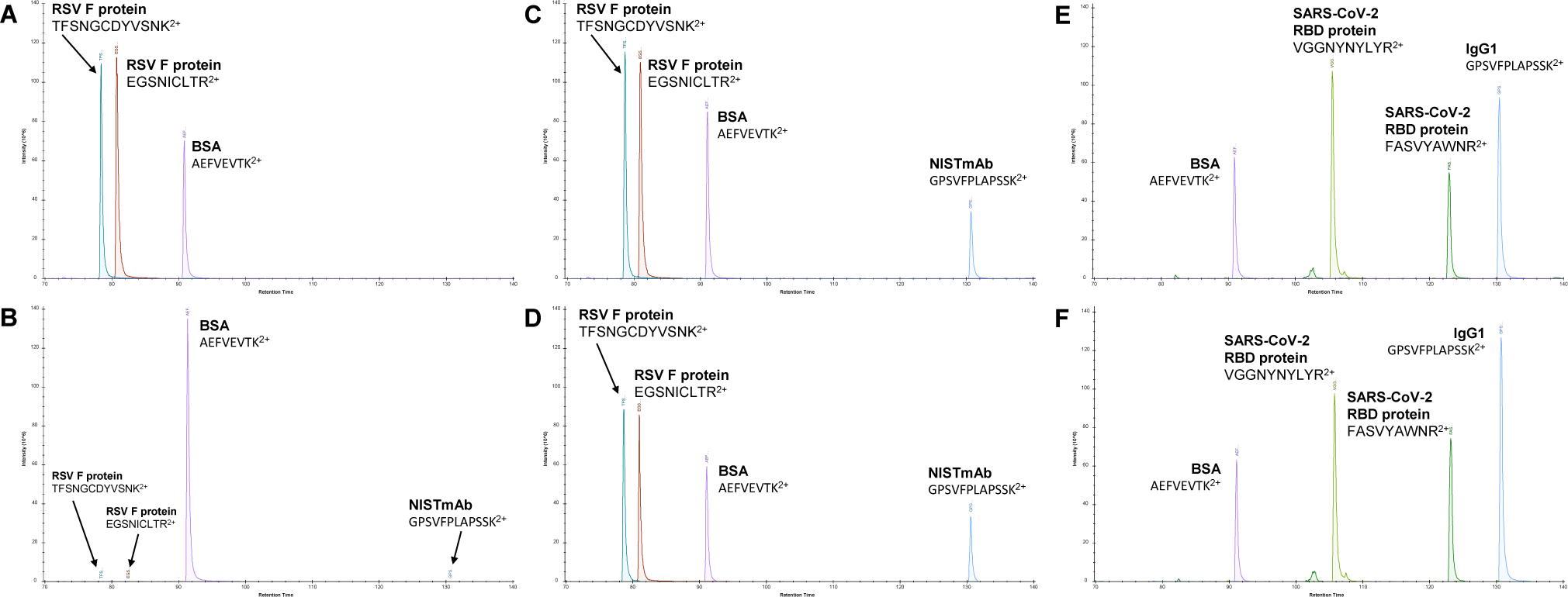
Enrichment of NISTmAb 8671 with an RSV preF protein, as measured by IA-MS assay with label-free MS1 quantification. **(A)** Positive control: evidence of preF (0.5 µg/well) coating onto immunoassay microplates. Two tryptic preF peptides were measured. BSA peptide was presented as a loading control. **(B)** Negative control: lack of non-specific binding of NISTmAb 8671 monoclonal antibody (1 µg/well) to the microplates blocked with BSA. **(C-D)** Two analytical replicates revealed that NISTmAb 8671 (1 µg/well) was enriched with preF protein (0.5 µg/well). **(E-F)** Positive control for IA-MS: two analytical replicates provided evidence that SARS-CoV-2 RBD protein (0.5 µg/well) enriched IgG1 subclass of human antibodies present in a 20 µL pool of convalescent sera. Two tryptic peptides of SARS-CoV-2 RBD protein (0.5 µg/well) were presented. NISTmAb 8671 antibody and the endogenous human IgG1 antibodies shared a unique tryptic peptide GPSVFPLAPSSK.

### Differential quantification of immunoglobulin isotypes and subclasses

Our approach to identification and quantification of polyclonal antibodies is presented in **Fig. 4** and was discussed in detail in our previous studies^14,18,41^. We previously demonstrated IA-MS as an experimental approach for rational design and development of SARS-CoV-2 serological diagnostics^14^. Briefly, 20 µl of healthy normal serum or plasma were subjected to IA enrichment onto microplates coated with preF protein. The purified synthetic heavy-isotope labeled peptide internal standards (300 fmoles/well; SpikeTides TQL) unique for each isotype and subclass were added prior to trypsin digestion and used for “absolute” quantification based on L/H ratio measured by nanoLC-MS, as previously reported^14^. Here, we measured absolute concentrations of anti-RSV preF antibody isotypes and subclasses in 69 serum samples of healthy individuals and revealed IgG1 (2,580 ng/mL), IgA1 (280 ng/mL) and IgM (180 ng/mL) as the most abundant isotypes **(Table 1**). High levels of IgG1 immunoglobulins were found in all samples, as compared to the assay’s negative controls **(Fig. 5)**. Interestingly, we also identified the statistically significant presence of IgG2 (74 ng/ml), IgG4 (4.9 ng/mL), and IgA2 (5.5 ng/mL) isotypes. The biological and clinical significance of these data must be further investigated.

**Fig 4.**
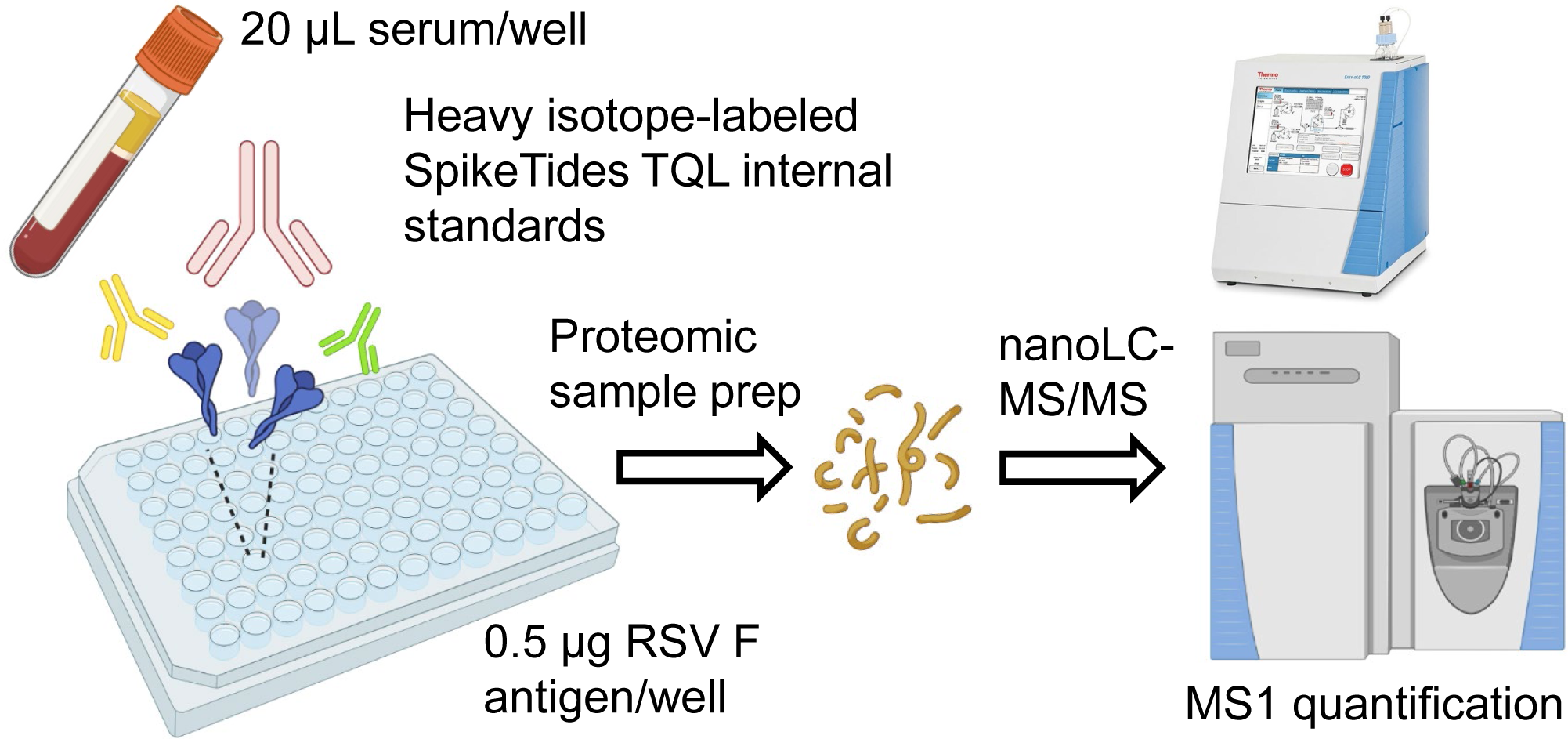
Experimental approach for differential quantification of immunoglobulin isotypes and subclasses.

**Table 1.** Serum concentrations of anti-RSV immunoglobulin isotypes and subclasses, as measured by IA-MS. (+) RSV group included 69 serum samples, while negative control group (–) PBS included 3 matched serum samples. Median concentrations and interquartile ranges [IQR] were presented. MWU, Mann-Whitney U test.

|  | IgG1 | IgG2 | IgG3 | IgG4 | IgM | IgA1 | IgA2 | IgD |
| --- | --- | --- | --- | --- | --- | --- | --- | --- |
| (+) RSV,<br>ng/mL | 2,580 | 74 | 5.1 | 4.9 | 177 | 277 | 5.5 | 5.2 |
| [IQR] | [1,820-4,200] | [51-122] | [2.7-11.9] | [3.7-10.1] | [102-354] | [163-563] | [4.3-11.2] | [5.2-6.5] |
| (–) PBS,<br>ng/mL | 160 | 25 | 2.5 | 3.7 | 51 | 86 | 4.3 | 5.2 |
| Fold Change | 16.1 | 3.0 | 2.0 | 1.3 | 3.4 | 3.2 | 1.3 | 1.0 |
| MWU *P*-value | 0.002 | 0.008 | 0.114 | 0.037 | 0.011 | 0.010 | 0.001 | 0.154 |

**Fig 5.**
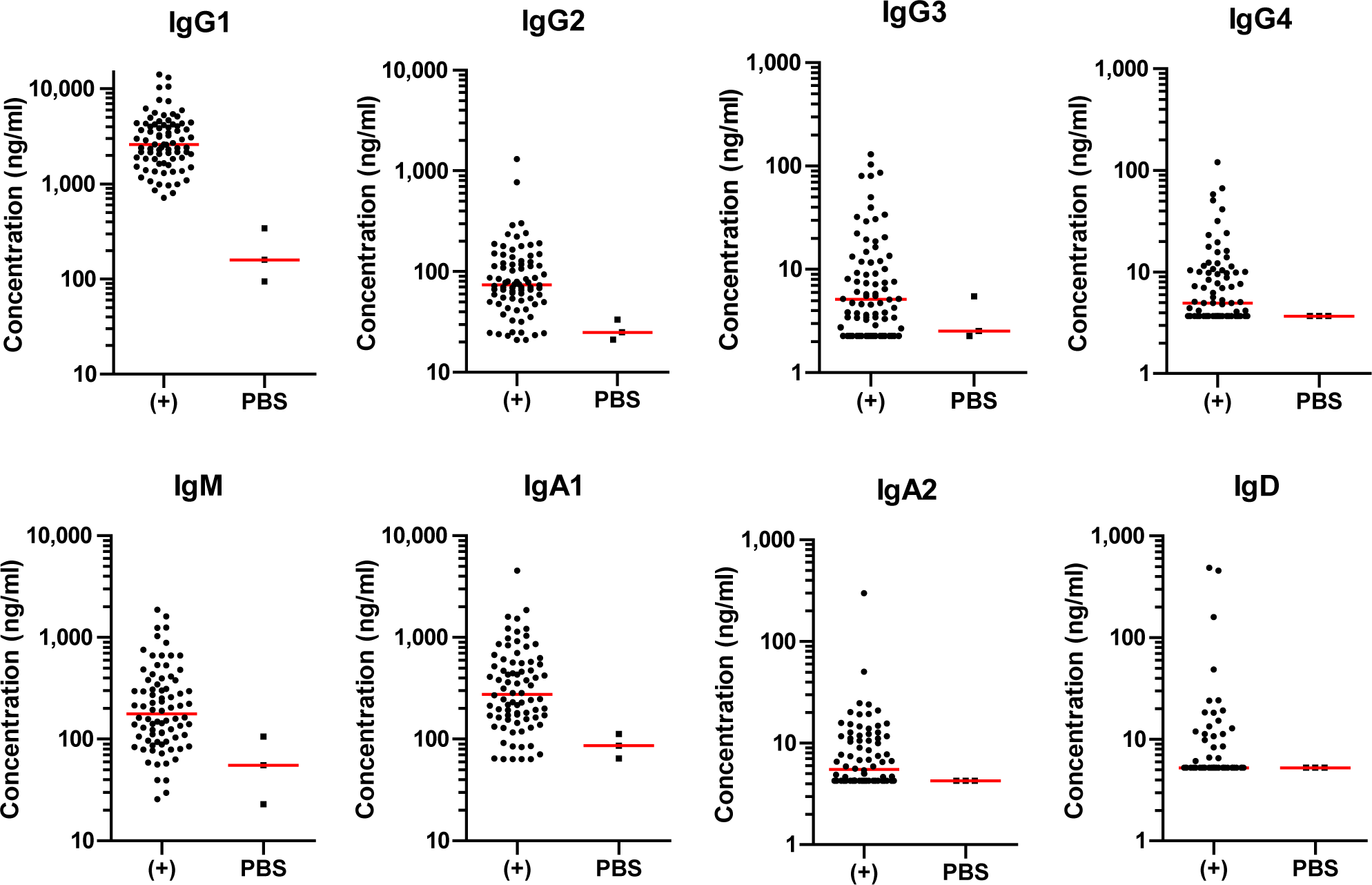
Differential quantification of immunoglobulin isotypes and subclasses.

### Co-enrichment of the endogenous Fc interactome

Anti-RSV antibodies were enriched with RSV preF protein (+RSV) and PBS (–PBS negative control) from 62 and 3 matched serum samples, respectively. The search of IA-MS data against the canonical UniProt human proteome revealed the consistent enrichment of the C1q complement complex proteins C1qA, C1qB, and C1qC, as well as CD5L protein (**Fig. 6**). Interactions of C1q complex with IgG1 and IgG3, as well as the interaction of CD5L with IgM have been well established^72,73^.

**Fig. 6.**
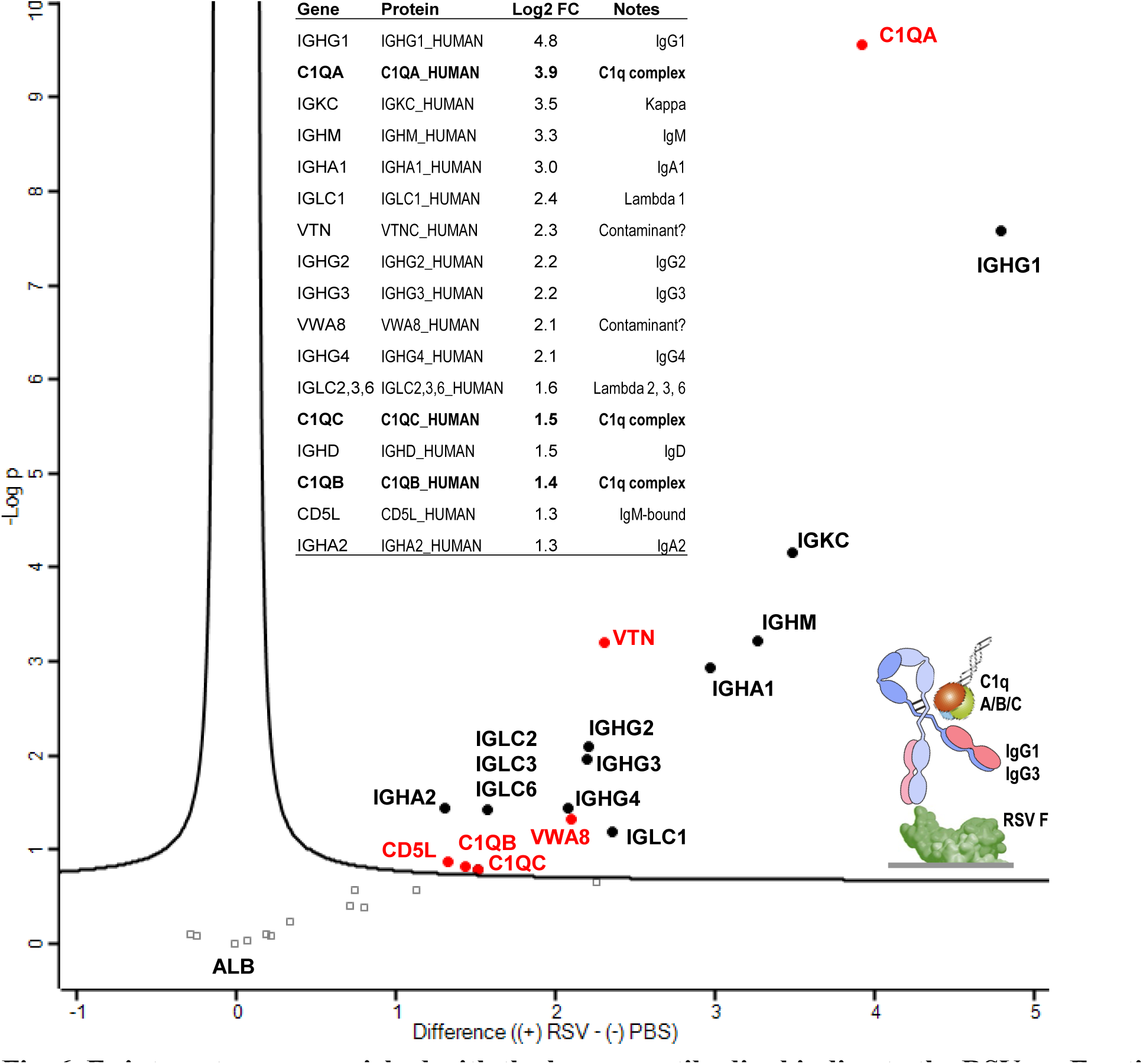
Fc interactome co-enriched with the human antibodies binding to the RSV preF antigen. The results revealed consistent enrichment of the C1q complement complex proteins C1qA, C1qB, and C1qC (potentially co-precipitated together with IgG1 and IgG3), as well as CD5L protein (potentially co-precipitated with IgM). Anti-RSV antibodies were enriched from 62 serum samples with RSV F antigen protein (+RSV) and 3 matched serum samples without an antigen (–PBS). PBS negative control: microplates coated with Phosphate-Buffered Saline instead of an antigen. Label-free quantification approach (MaxQuant LFQ values) was utilized for the quantitative measurements. The table presents gene and protein names, and corresponding log2-transformed fold changes (log2 FC).

### IGHV gene usage

Polyclonal antibodies are highly diverse in terms of the usage of the germline variable (V), diversity (D), joining (J), and constant (C) genes. According to the IMGT database^74^, the germline sequences of the human antibody heavy chains are composed of one of 56 functional IGHV genes (with 289 alleles), one of 24 functional IGHD genes (30 alleles), one of 6 IGHJ genes (13 alleles), and one of nine IGHC genes (such as IGHC1 for the IgG1 subclass). Likewise, the germline sequences of the human light kappa chains are composed of one of 39 functional IGKV genes (69 alleles), one of 5 functional IGKJ genes (6 alleles), and an IGKC gene (kappa light chains). The germline sequences of the human light lambda chains are composed of one of 32 functional IGLV genes (81 alleles), one of 7 functional IGLJ genes (7 alleles), and one of 5 functional IGLC genes (such as IGLC1 for lambda 1 light chains). Somatic hypermutation and affinity maturation introduce point mutations across V(D)J variable regions and substantially modify CDR1, CDR2, and CDR3 sequences, with CDR3 regions being the most variable and hardly resembling the initial germline sequences^75^. The complete sequencing of the functional antibody repertoire by single-cell RNA sequencing of memory B cells and plasmablasts has recently been established^75,76^. Numerous studies revealed that polyclonal antibody response was dominated by only a few most frequent germline genes (e.g., IGHV3-23)^77^.

Here, we explored the hypothesis that IA-MS could emerge as a viable approach to sequence the functional antibody repertoires. We focused on IGHV germline genes that define the bulk of VH antibody scaffolds (∼100 aminoacids) and may include numerous tryptic peptides suitable for LC-MS/MS identification and relative quantification. To search for IGHV genes, we generated an in-house database that included the complete UniProt human proteome, 44 most abundant IGHV genes based on their occurrence in the human B cell transcriptome (>0.5% frequency in the population), and 7,272 IGHV sequences of the human antibodies derived by single-cell RNA sequencing of human B cells^78^. We acknowledge that our current database could be incomplete and could miss some high-abundance peptides with point mutations.

The MaxQuant search of IA-MS data for 69 individuals resulted in the identification of 400 unique IGHV protein sequences (at a 1% false discovery rate), 556 unique IGHV peptides, and the corresponding 2,771 MS/MS spectra. In this population, we identified 5 IGHV subgroups, with IGHV3 subgroup being the most abundant (**Fig. 7A**). In total, we identified in our population the expression of 47 unique IGHV genes, with 90% and 50% of abundances represented by 22 and 7 unique genes, respectively (**Fig. 7B**). The sample with the highest diversity of IGHV repertoires (E2) revealed 31 unique IGHV genes, with 90% and 50% of abundances represented by 25 and 8 unique genes, respectively. The sample with the lowest diversity of repertoires (E1) revealed 18 unique IGHV genes, with 90% and 50% of abundances represented by 8 and 1 unique IGHV genes, respectively (**Fig. 7B**). While the biological significance of these data must be further investigated, the presented study paves the way to detailed characterization of the circulating antibody repertoires directly from the serum samples.

**Fig. 7.**
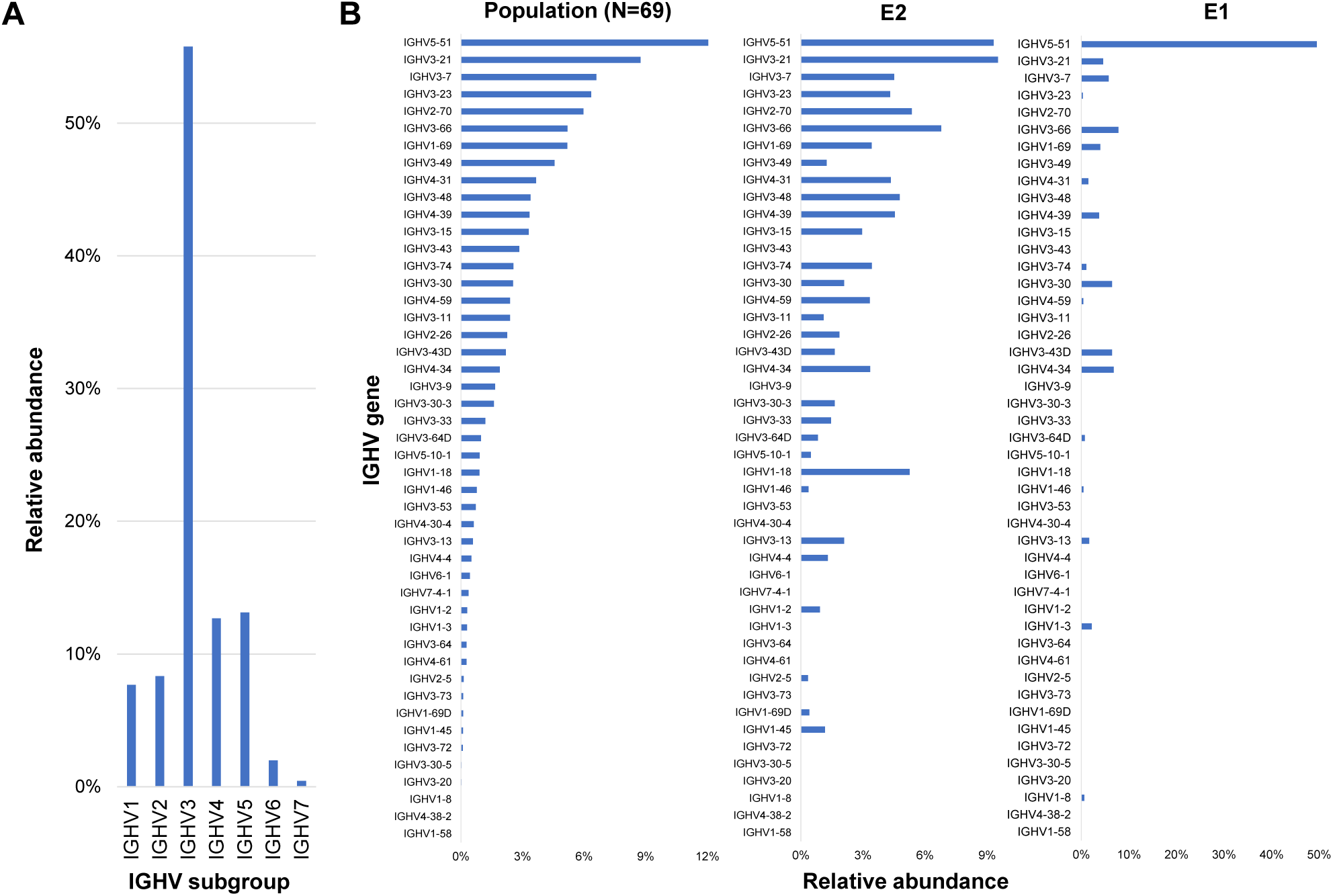
IGHV repertoire profiling of polyclonal antibodies against RSV preF protein. Serum samples of 69 individuals were subjected to IA enrichment, proteomic sample preparation, and LC-MS/MS measurements. MaxQuant iBAQ values for each IGHV gene identification were normalized by IgG1 iBAQ values. Relative abundance was calculated as the sum of normalized iBAQ values for each subgroup over the total normalized iBAQ signal. **(A)** Subgroup IGHV3 was revealed as the most abundant (56%). **(B)** Relative abundance of IGHV gene usage in population (N=69 samples), the sample with the highest diversity of repertoires (E2), and the sample with the lowest diversity of repertoires (E1) based on our data. While 23 genes contributed to the use of the IGHV3 subgroup, the most abundant unique gene was IGHV5-51.

## CONCLUSIONS

We presented a proteomic toolbox for the identification, quantification, and characterization of the endogenous polyclonal antibodies. We demonstrated our approaches with the human antibodies generated against RSV, but the proposed methods could be extended to the investigation of animal antibodies and the numerous antigens of infectious diseases and non-infectious disorders^41^. The presented toolbox will facilitate the in-depth characterization of polyclonal antibodies and pave the way to quantitative approaches in serological testing and precision immunology.

## Acknowledgements

We thank Yasmine Rais for guidance and assistance with proteomic sample preparation, and Yuan Yuan Zhao for assistance with maintenance of Orbitrap Elite mass spectrometer.

## Author Contributions

A.P.D. designed the research project; W.T. performed the experiments; A.P.D. and W.T. analyzed data; A.P.D. wrote the manuscript, and all authors contributed to the revisions.

## Non-standard abbreviations

ELISA: enzyme-linked immunosorbent assay
iBAQ: intensity-based absolute quantification
IA-MS: immunoaffinity-mass spectrometry
LC-MS/MS: liquid chromatography - tandem mass spectrometry
LFQ: label-free quantification
MWU: Mann Whitney Unpaired t-test
RSV: respiratory syncytial virus.

